# Brain Functional Connectivity Correlates of Anomalous Interaction Between Sensorily Isolated Monozygotic Twins

**DOI:** 10.1101/2023.10.18.563012

**Authors:** Richard B. Silberstein, Felicity J. Bigelow

## Abstract

This study examined brain functional connectivity (FC) changes associated with possible anomalous interactions between sensorily isolated monozygotic twins. Brain FC was estimated using the Steady State Visual Evoked Potential-Event Related Partial Coherence (SSVEP-ERPC) methodology. Five twin pairs served twice as participants with an average interval between sessions of 67 days. In each recording session, one twin, the sender, viewed a randomized set of 50 general images and 50 personally relevant images, whilst the other twin, the Receiver, viewed a static personally relevant image for the entire duration of the session. Images appeared on the sender screen for 1.0 sec with the interval between successive images varied randomly between 4.0 and 8.0 sec. Receiver FC changes were calculated from the appearance times of the images as viewed by the Sender. It was hypothesized that anomalous interactions would be indicated by statistically significant Receiver FC changes when those changes are determined using the sender image appearance times. For each twin serving as Receiver, FC components were separately analysed for the 50 general and the 50 personal images, yielding 38 observations (19 twin pairs by 2 conditions). The hypothesis was confirmed in that 12 of the 38 observations, yielded statistically significant receiver FC increases or decreases at the p<0.01 level only when trials were synchronized to the Sender image appearance times. Overall, this effect was significant at the p=4X10^-8^ Df=175. To the best of our knowledge, this is the first study reporting statistically significant FC changes indicative of anomalous interactions between two sensorily isolated individuals.

## Introduction

Identical (monozygotic) twins have long been a topic of popular interest, if not intrigue. One aspect of such interest are claims that identical twins are more likely to report anomalous ‘shared experiences’ (Brusewitz et al 2013). An anomalous shared experience can be defined as the sharing or communication (through information, images, or emotions) between separate minds (Radin et al 2004, 2017). Typical of such anomalous ‘shared experiences’ are claims that one twin experienced somatic pain or discomfort at the same time their sibling was involved in a mishap or accident, even though the twins may be separated by a long distance (Brusewitz et al 2013).

A number of studies have examined experimental evidence for electroencephalographic (EEG) or functional Magnetic Resonance Imaging (fMRI) indications of correlated brain activity in pairs of individuals sharing an emotional bond such as a family relationship, friendship or a romantic relationship (Achterberg et al 2005, Brusewitz et al 2013, Duane & Behrendt 1965, Parker & Jensen 2013, Standish et al 2003, 2004). Typically, simultaneous brain activity is recorded from a pair of sensorily isolated individuals and one of the pair, ‘the Sender’ is exposed to a stimulus or a task. Anomalous interactions between the individuals are thought to be indicated by concurrent brain activity changes in the other participant, the ‘Receiver’. One of the earliest EEG studies of anomalous interaction in sensorily isolated twins involved the ‘Sender’ closing their eyes at various times to increase EEG alpha activity (Duane & Behrendt 1965). A concurrent EEG alpha activity increase in the receiver twin was considered evidence of anomalous interaction, with this effect reported in 2 of the 15 pairs of twins examined. A more recent fMRI study of a pair of monozygotic twins explored changes in brain activity during auditory and visual tasks (Karavasilis et al 2018). One twin was in the fMRI scanner, while the other twin outside the scanner passively viewed either emotional and personally relevant images provided by the twins, or heard sounds chosen to elicit fear. Both the visual and auditory task presentations were 260 sec in duration and comprised 20 sec stimuli followed by 20 sec rest intervals. Findings illustrated statistically significant increases in brain activity at left orbitofrontal gyrus, left cingulum, and left precentral gyrus at times the other twin was engaged in the auditory or visual perception tasks. Together, these studies provide support for the reality of such anomalous interactions.

While the majority of studies have reported evidence consistent with enhanced correlated brain activity in pairs of emotionally related individuals (Achterberg et al 2005, Karavasilis et al 2018, Radin 2004, Standish et al 2003, 2004, Wackermann et al 2003, Wackermann 2004), others have either failed to replicate these findings (Ambach 2009, Moulton & Kosslyn 2008) or have been criticized on the basis of small numbers of participants (Storm & Tresoldi 2017). Primarily, these studies have implemented tasks with little or no emotionally relevant stimuli, such as Zener cards (Billows & Storm, 2016; Bouvet & Bonnefon, 2015). However, past research has demonstrated that anomalous shared experiences tend to be personally relevant and emotional, rather than neutral (Brusewitz et al 2013; Radin & Schlitz, 2005). Therefore, is it theorized that the inclusion of emotional stimuli deemed personally relevant may yield a stronger effect.

While anomalous interbrain interactions (AII) have been reported between individuals sharing a strong emotional bond, we note that such effects have been more commonly reported in monozygotic twins (Brusewitz et al 2013, Parker & Jensen 2013, Radin 2004). As we wished to design this study in a way to maximize our chances of observing AII we restricted the participant recruitment to monozygotic twins. Importantly, the decision to restrict recruitment to monozygotic twins does not represent any attempt to compare findings from monozygotic twins with any other pairs of participants.

To date, the two main brain imaging methodologies that have been used to study inter brain anomalous interactions have been EEG and fMRI. The current study breaks new ground in being the first study to examine dynamic changes in brain functional connectivity (FC) that may indicate the presence of AII. This approach is novel in applying the evoked potential methodology Steady State Visual Evoked Potential Event Related Partial Coherence (SSVEP-ERPC) to measure brain FC changes indicative of AII.

The choice of SSVEP-ERPC was based on several factors. Firstly, unlike fMRI, the methodology offers relatively high temporal resolution estimates of brain FC enabling one to identify cognitive task components associated with specific FC changes (Silberstein et al 2003). Additionally, the SSVEP-ERPC is highly resistant to the most common sources of EEG noise and artifacts (Gray et al 2003). Finally, the methodology has been found to be especially sensitive to FC changes associated with cognitive processes, such as the attentional and hyperactivity components of ADHD (Silberstein et al 2016A, 2016B, 2017). The technology has also contributed novel findings concerning the dynamic features of brain connectivity networks that mediate the level of originality in creative cognition (Silberstein et al 2019).

In the current study, changes in brain FC were examined in five pairs of monozygotic twin females, comprising a series of single pair (Sender and Receiver) studies. In this paper the twin viewing the images will be referred to as the ‘Sender’ and the other as the ‘Receiver’. Fifty of the images portrayed landscapes (non-personal images) while the remaining 50 were images provided by each set of twins and were selected as being of personal or emotional relevance to the twins (personal images). The critical metric we will be evaluating is the number of simultaneous statistically significant event FC changes in the Receiver when we use the image presentation times as viewed by the Sender. This is the metric that has been used in previous studies examining the effects of cognitive processes on brain FC and has revealed evidence of both FC increases and decreases associated with cognitive task performance (Silberstein et al 2016A, 2016B, 2017, 2019). As this study constitutes the first application of the SSVEP-ERPC methodology to examine AII, our hypotheses are restricted to the predicted observation of FC changes rather than the direction or topography of those FC changes.

### Hypothesis 1

The presence of anomalous interactions between monozygotic twin pairs will be indicated by a statistically significant number of simultaneous event-related FC changes in the Receiver only when the receiver event related FC is determined using the image presentation times as seen by the Sender.

### Hypothesis 2

In the event that statistically significant Receiver FC changes are observed, these will be larger for the cases where the Sender is viewing personal images.

## Material and methods

### Ethics approval and informed consent

Informed consent was obtained from all participants, and from guardians of participants aged under 18 years. Ethics approval for the study was obtained from Swinburne University of Technology Human Ethics Research Board (# SHR Project 2018/416).

### Scheduling

All twins participated in two recording sessions. Recording sessions took place in Nov 2022 (Session 1) and Jan-Feb 2023 (Session 2).

Each participant acted as Sender and Receiver in the recording sessions, with participants randomly assigned roles.

### Participants

To be eligible for participation, individuals were required to be monozygotic twins aged 13 years or older. Monozygosity was determined through DNA testing that was made available to all twins participating in the study. Overall, the sample consisted of 5 pairs of female monozygotic twins. The results of monozygosity testing and age of participants is provided below.

In addition to monozygosity, other selection criteria were, a belief held by both twins that they had previously experienced at least one shared anomalous experience and that both twins were able to attend the same recording session.

Exclusion criteria included, pregnancy, epilepsy, a history of head trauma, or a current or past diagnosis of schizophrenia or dissociative disorder.

### Viewing content

Images presented during the recording tasks included 50 personally relevant and 50 general images. All images were presented on a black background and subtended a vertical angle of 26 degrees and a horizontal angle of 14 – 26 degrees depending on the image. Each participating pair was asked to provide no more than 50 and no less than 25 personally relevant images. Personally relevant images were defined as photographs representing something significant or meaningful to the twin pair. Where necessary, personally relevant images were randomly repeated to reach 50 trials. All personally relevant images provided by participants were treated with confidentiality.

Fifty general images were sourced from a free online database. These images included landscape and natural scenery stimuli. Each participant viewed the same set of general images but in a unique random order.

### Procedure

Participants attended two 90-minute sessions roughly two months apart (*M _interval_* = 67 days). Each session included recording tasks, followed by a series of questionnaires.

### Session 1 Procedure

After providing informed consent, participants were randomly and blindly assigned the role of Sender or Receiver. Participants were then seated in separate rooms roughly eight meters apart. Rooms did not share any walls and were further isolated by three closed doors.

Participants were seated approximately 60 centimeters from a 32-inch screen, in a quiet dark room. Participants were informed that they would be presented with one or more images on a screen and to passively view the image or images. During the recordings in both Sessions 1 and 2, twins were seated in separate rooms separated by approximately 8 m and three closed doors. Data acquisition and image presentation was controlled by two computers, each computer located near each twin. Fig 1 illustrates the relative locations of the twins and data acquisition equipment during recording sessions 1 and 2.

**Fig 1.**
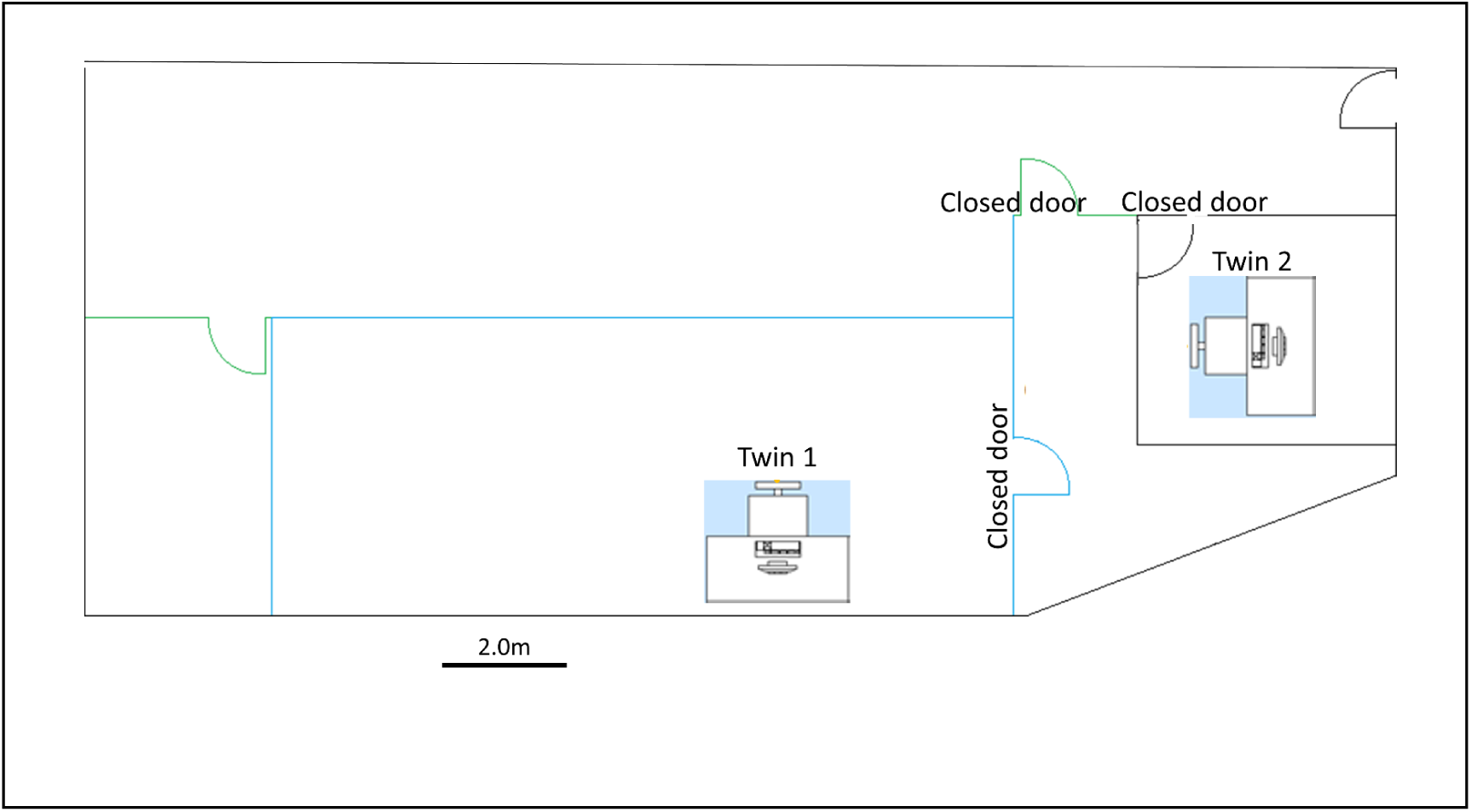
Recording rooms arrangement and location of participants during recording.

During Session 1, twin pairs completed two recording tasks, serving successively as Sender and Receiver or in the reverse order. Following the first task, participants spent approximately 5 minutes swapping both rooms and roles.

#### Sender

The Sender was presented with 100 visual stimuli comprising both general and personally relevant images. Each image appeared on the screen for 1.0 second. The appearance of each image was followed by a 4.0 second fixed interval and a unique random interval 0.0 second – 4.0 second determined by a TrueRNG random number generator (*M _interval_* = 6.0 sec; *SD* = 1.2 sec). The order of the 100 images was randomized for each twin using the Matlab Randperm function, with the task lasting roughly 11 minutes. While the average interval between successive images was 6 sec (SD=1.2 sec), the average interval between successive general or personal images was 12 sec (SD=2.4 sec).

#### Receiver

The Receiver was presented with one personally relevant visual stimulus depicting both twins. The presentation of this image remained for the entire task, lasting approximately 11 minutes.

Each participant was reimbursed $200 for their time in Session 1.

### Session 2 Procedure

Session 2 followed the same procedure as Session 1 recording with the following two differences. The two image sequences viewed by the Senders in Session 1 were swapped in Session 2. In other words, the image sequence viewed by Sender 1 in Session 1 were now viewed by Sender 2 in Session 2 and vice versa. The second difference was the inclusion of an additional recording component at the end of the sender-receiver recordings. This was an 11-minute control recording where both twins viewed a single personal image. During this time, twins were instructed to passively view the image. This final task acted as an additional control component.

Each participant was reimbursed $200 for their time in Session 2.

### Data acquisition and the Steady State Visual Evoked Potential (SSVEP)

Brain electrical activity was recorded from 20 scalp sites in line with the International 10-10 system (Acharya et al 2016). The average potential of both mastoids served as a reference, while an electrode located at FPz served as a ground. Brain electrical activity was amplified and bandpass filtered (3 dB down at 0.1 Hz and 30 Hz) before digitization to 16-bit accuracy at a rate of 400 Hz. The major features of the signal processing have previously been described (for further details see Silberstein et al 2016A).

The stimulus used to evoke the SSVEP was a spatially diffuse 13-Hz sinusoidal flicker subtending a horizontal angle of 160° and a vertical angle of 90°, which was superimposed on the visual fields. This flicker was present throughout the task and special goggles enabled subjects to simultaneously view the cognitive task and the sinusoidal flicker. The modulation depth of the stimulus when viewed against the background was 45%.

A three-step process was used to calculate the steady state visual evoked potential (SSVEP). The first step involved band-pass filtering the sampled data (11.0 Hz – 15.0 Hz) using the fast Fourier transform. For the second step, the discrete Fourier transform was used to determine the real and imaginary single cycle Fourier coefficients at the 13Hz stimulus frequency. The final step involved smoothing the real and imaginary Fourier coefficients with a cosine weighted smoothing window 23 stimulus cycles in length. The equivalent or rectangular window width is approximately half the duration of the cosine width or 12 cycles. The cosine smoothing window was then shifted 1 stimulus cycle, with the coefficients recalculated for this overlapping period. This produced the SSVEP and was continued until the entire 380 seconds of activity was analyzed. An identical procedure was applied to data recorded from all recording sites.

### SSVEP Event-Related Partial Coherence (SSVEP-ERPC)

Functional connectivity (FC) between electrode sites was determined using an approach termed SSVEP Event Related Partial Coherence (SSVEP-ERPC). The SSVEP-ERPC is a measure of the partial coherence between electrode pairs at the stimulus frequency eliciting the SSVEP and is based on a modification of an approach first described by Andrew and Pfurtscheller (1996) (Silberstein et al 2003).

Partial coherence varies between 0 and 1 and, like coherence, is a normalized quantity that is not determined by the SSVEP amplitude at either electrode site. Electrode pairs with high partial coherence indicate relatively stable SSVEP phase differences between electrode pairs across trials. This occurs even though SSVEP phase differences between each of the electrodes and the stimulus may be variable across trials and is equivalent to the removal of the common contribution from the SSVEP stimulus. Therefore, the high SSVEP partial coherence between electrodes reflects a consistent synchronization between electrodes at the stimulus frequency, rather than a consequence of two unrelated regions increasing their response to the common visual flicker. Such synchronization reflected in the SSVEP-ERPC is thought to reflect FC between the relevant regions, with the terms ‘SSVEP-ERPC’ and FC used interchangeably.

For each Sender and Receiver, the SSVEP-ERPC was calculated for all 190 distinct pairs of electrodes averaged for both the personal image presentation times and the general image presentation times <u>as viewed by the sender</u> in each of the recording sessions. In both cases, personal and general images, the SSVEP-ERPC was calculated for a 5.0 sec epoch commencing 2.0 sec before the appearance of an image.

### Statistical analysis

Hypothesis 1 is based on the assumption that the presence of anomalous interactions between the twins will be indicated by the number of simultaneous statistically significant FC changes (out of 190) in the receiver when those observed FC changes are estimated using the image presentation times <u>as viewed by the sender</u>.

A four-stage process was used to identify the observed number of statistically significant FC increases and decreases at each point in time. Each of the four stages are outlined in Fig. 2 and then described in more detail subsequently. For a more general discussion on the application of nonparametric statistical testing of coherence differences, the reader is referred to Schoffelen & Fries, (2007).

**Fig 2.**
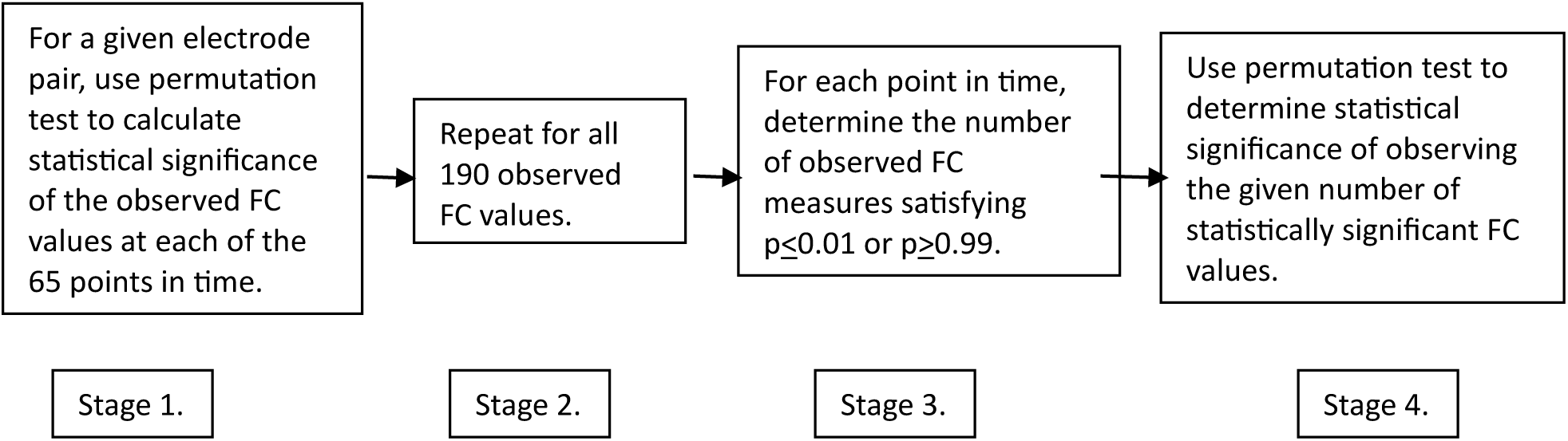
Discrete stages in the statistical analysis.

### Identifying statistically significant FC changes

To illustrate Stage 1 of the procedure, we first consider a simplified case comprising one electrode pair (F_z_ – P_3_) in one subject, in this case acting as the sender, that is the twin viewing the images. Fig 3 illustrates the 5-sec (F_z_-P_3_) event related FC time series calculated on the appearance times of the <u>general images</u>. We assume that the specific FC maxima and minima emerge only when we use the actual image presentation times.

**Fig 3.**
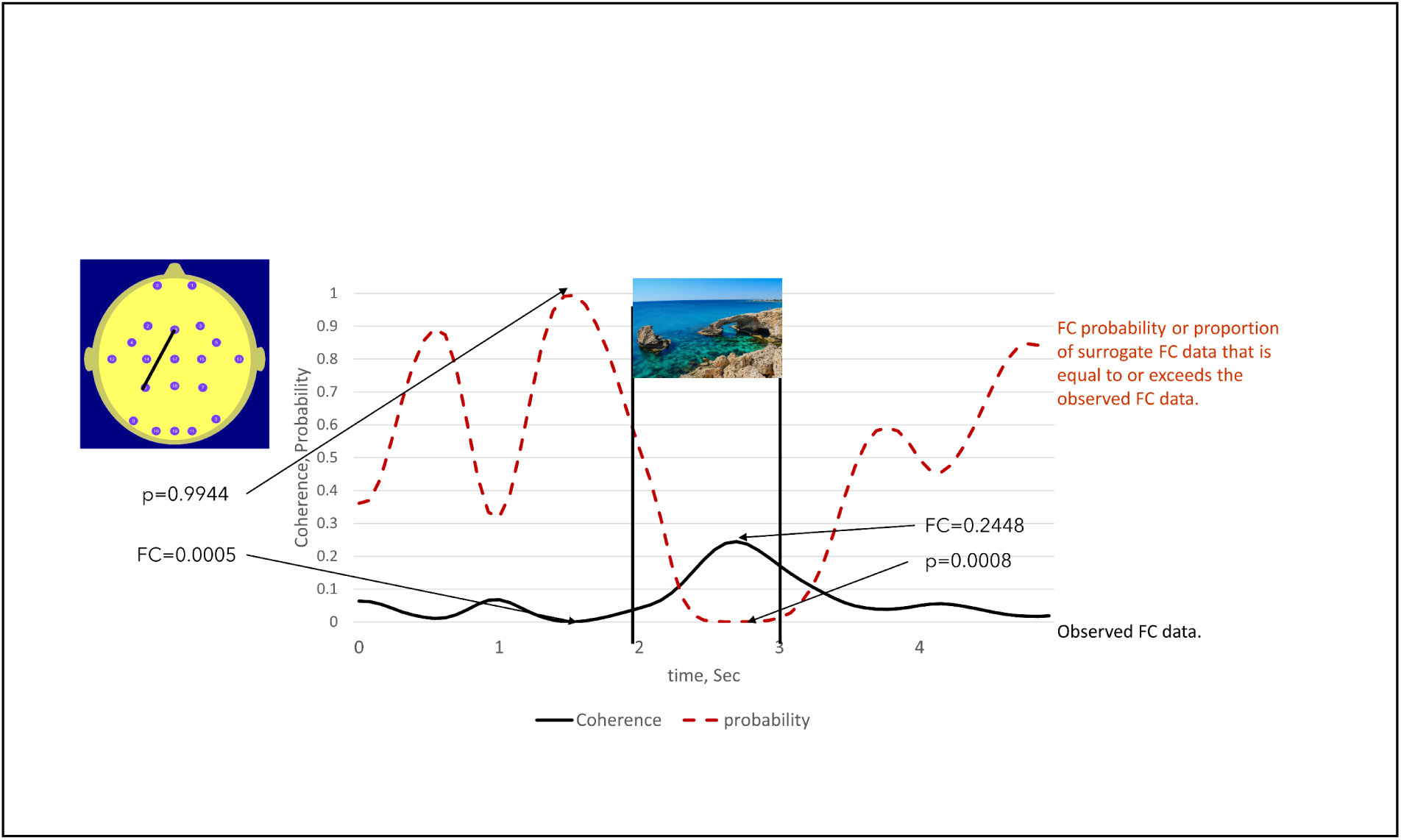
illustrates the use of the permutation test to identify statistically significant high and low FC values in a participant viewing the images (Sender). The solid line is the observed FC between electrodes Fz – P3 calculated on the appearance times of 50 general images, here represented by a sea scene. The dotted line indicated the proportion of times that the surrogate FC value equalled or exceeded the observed FC value. This was equivalent to the probability of the peak FC value being observed when image presentation times are used to calculate the observed FC. The statistical significance of observing low FC values can also be determined using this approach. The low observed FC value at 1.5 sec is exceeded by 99.44% of the surrogate FC values. Thus, the probability of observing such a low (as opposed to a high) FC value is thus: p=100% -99.44% = 0.56%.

In this case, the NULL hypothesis states that the observed FC maxima and minima are random events and unrelated to the image presentation times. To test the NULL hypothesis, we created a random array of image presentation times with the same mean inter-image interval and recalculated the event related FC, subsequently referred to as the surrogate FC. This step is repeated 10,000 times and the number of times the observed FC values are exceeded by the surrogate FC values is noted.

The proportion of the 10,000 surrogate FC values that exceeded the observed FC values at any specific point in time, is then the probability (p) that an observed FC maxima at that point in time could have occurred by chance. If p is less than a threshold value, typically 0.05 or 0.01 then the NULL hypothesis can be rejected at this level of a likely type 1 statistical error. We can also use this approach to determine the statistical significance of observed FC minima. For example, if 99% of the surrogate FC values exceed an observed FC minimum, then that minimum value is statistically significant at the 1% level (1-p).

Stage 2 involved repeating the Stage 1 process for all 190 FC measures and for all points in time in the 5.0 sec (65 point) epoch.

### Accounting for multiple comparisons

To account for the multiple comparisons in determining the statistical significance of the FC value at each of the 65 points in time, a threshold value of p≤ 0.01 rather than p≤0.05 was selected as the criterion for statistical significance. Adjustments for multiple comparisons are most commonly achieved using a Bonferroni correction, where the nominal level for statistical significance, typically p=0.05 is divided by the number of independent observations or degrees of freedom. However, the 65 points of the FC time series are not statistically independent, with neighbouring points being highly correlated primarily due to the cosine smoothing window as well as the Fourier filtering at an earlier stage of data analysis.

To determine the effective number of degrees of freedom of the FC time series, we first estimate a parameter known as the e-folding decay time (e-fold). The e-folding decay time is the interval over which the autocovariance of the smoothed SSVEP time series falls to 1/e of its value at the zero interval. (Panofsky and Brier, 1958, Yu, J.-Y. 2005). Equation 1 relates the e-fold to the effective number degrees of freedom.

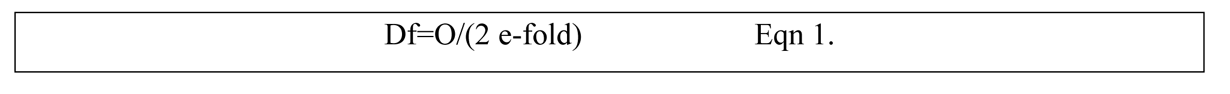

Where Df is the number of degrees of freedom, O is the number of observations, 65 and e-fold is the number of cycles over which the autocovariance decays to 1/e of its maximum value. The e-folding time in cycles for all electrodes was found to be 7 cycles. In this case, the number of degrees of freedom for each 65-point time series is 65/(2*7) or 4.64. By taking p≤0.05 as the threshold for statistical significance of the permutation test, then the p-value adjusted for the multiple comparisons was p=0.05/4.65 or p≤ 0.0108, which was rounded down to p≤0.01.

### Number of significant individual FC results (NSFC)

The next stage of the individual statistical analysis involved determining the number of observed FC measures out of 190 that were each simultaneously significant at the p_i_≤0.01 (FC maxima) or p_i_≥0.99 (FC minima) (Fig 2, Stage 3, NSFC). A permutation test was then used to determine the statistical significance of observing a given <u>number of FC measures</u> each satisfying the condition p_i_≤0.01or p_i_≥0.99 (Fig 2, Stage 4). This was determined separately for the 50 personal image times and the 50 general image times as viewed by the Sender.

An important point to note is that the metric under consideration, NSFC or the number of FC measures satisfying the statistical criteria, is a single positive integer defined at each of the 65 points in time. Thus, even though NSFC is determined from a consideration of all 190 unique FC measures, there is only one statistical test conducted at each of the 65 points in time. Once again, the Bonferroni correction is applied to the p<0.05 allowing for 4.65 independent measures or degrees of freedom.

To test the NULL hypothesis that the observed number of FC values satisfying the above p_i_ criteria (maxima and minima) could occur by chance, an array of random presentation times using the same mean inter-image interval of 12 sec were generated.

For each permutation, the number of surrogate FC maxima and FC minima that satisfied the conditions p_i_≤0.01 or p_i_≥0.99 were determined. This was repeated 10,000 times and the proportion of times that the number of surrogate FC measures satisfying the conditions of p_i_≤0.01 or p_i_≥0.99 was taken as the exact probability (p_g_) of observing this number of FC maxima or minima by chance (Fig 2, Stage 4).

For each individual, this process was repeated for the alternative image class, in this case, personal images. In other words, each participant as Receiver yielded two sets of results, one for each image class, (general images and personal images). These results will be referred to as *individual receiver findings* (IRF).

### Statistical significance of experiment wide results

To determine the statistical significance of the experiment wide results, we used the trinomial distribution, an extension of the binomial distribution to assess the probability of observing a given number of events where the ‘events’ are cases where the number of significant FC changes (NFSC) in the individual receiver findings satisfy the criterion p<0.01.

Each participant yielded two IRFs in each of the recording sessions. One individual Receiver finding for the Sender viewing general images and one for the Sender viewing personal images. In total there are N individual Receiver findings where N is twice the number of Receiver participant. In considering all N individual Receiver findings we designate n_1_ to indicate the number times that NFSC satisfies the condition 0.005<p≤0.01 and n_2_ to be the number of times that NSFC satisfies the condition p≤0.005. To determine the probability of observing, the n_1_ cases satisfying 0.005<p≤0.01and the n_2_ cases satisfying p≤0.005 a trinomial distribution with (N x 4.65) degrees of freedom was used. In using the trinomial distribution, we set the p values in the range 0.005<p≤0.01 to the value p_1_= 0.01 and the p values in the range p<0.005 to the value p_2_=0.005 in the following equation:

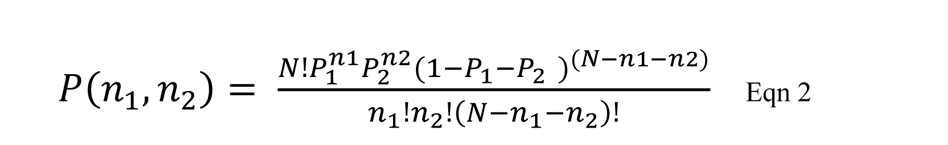

Where P(n_1_,n_2_) is the probability of observing n_1_ events with probability p_1_ and n_2_ events with probability p_2_ and (N-n_1_-n_2_) events with a probability of (1-p_1_-p_2_) (ie, non-significant observations).

The total number of independent observations being the number of individual receiver findings multiplied by the number of degrees of each individual observation or 4.65. It should be noted that this approach explicitly accounts for the total number of independent observations and hence accounted for these multiple observations.

## Results

Whilst the principal focus of this report was exploring the changes in Receiver FC, we illustrate the presentation format with an example of FC changes in a <u>Sender participant</u>.

Fig 4 illustrates the topography and number of statistically significant FC measures (NSFC) in the participant Sender 1_1 when viewing the general and personal images. The NSFC that are individually significant at the p≤0.01 and p≥0.99 level was indicated by the red and blue time series respectively. The statistical significance of the number of FC measures satisfying this criterion was indicated by the ‘p’ value above the maps. The p value applies to whichever parameter exceeds the threshold NSFC value corresponding to p<0.01. At the 0.8 sec mark the number of FC reductions (blue arcs) were statistically significant while at the 2.8 sec mark, the number of high FC components (red arcs) were statistically significant.

**Fig 4A.**
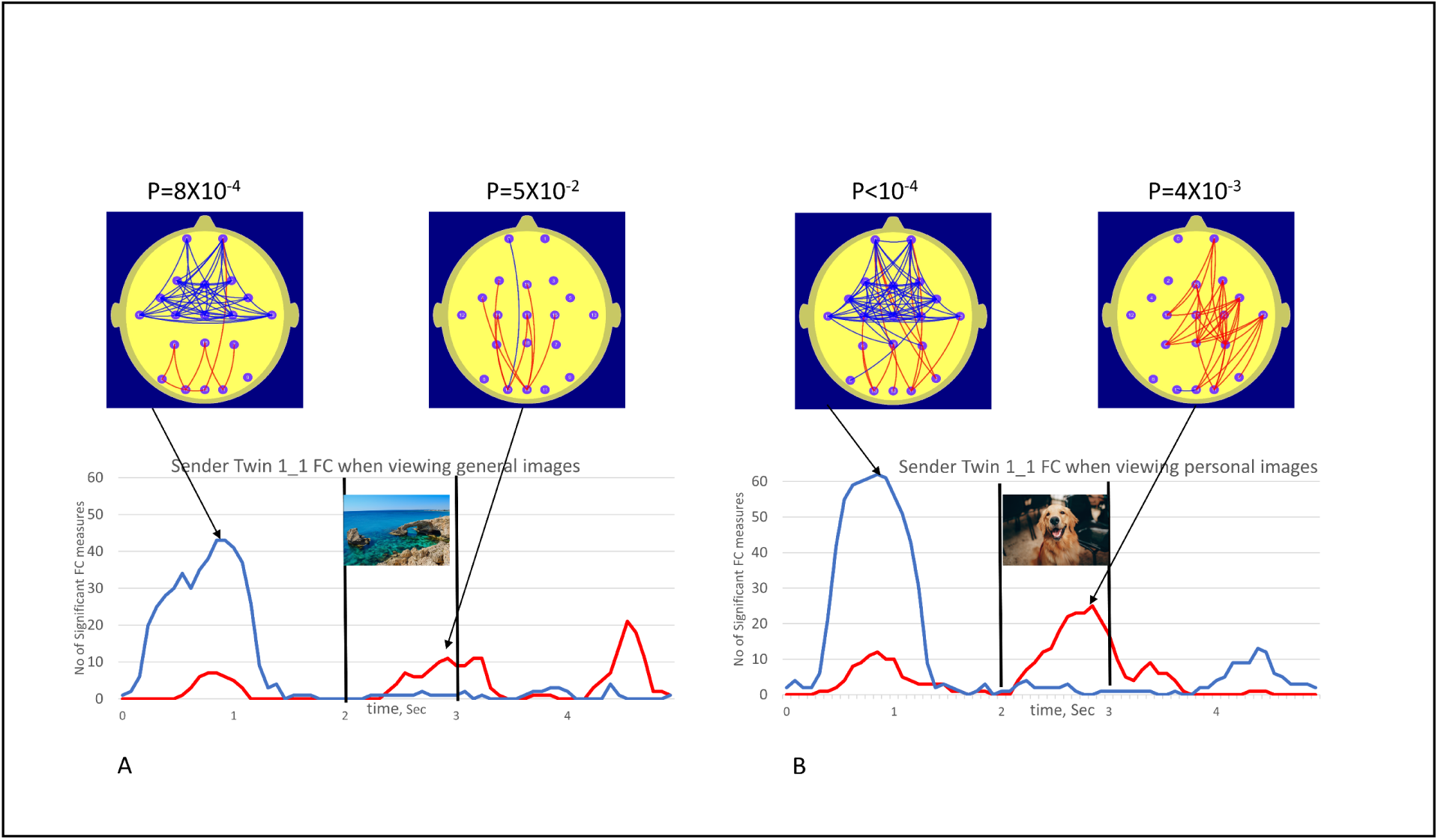
(left plots) illustrated the number of FC measures (NSFC) satisfying conditions p<0.01 (red trace, high FC values) and p>0.99 (blue trace, low FC values) in Sender 1_1 FC when viewing the 50 general images. Fig 4B (right plots) illustrated the equivalent for Sender 1_1 viewing personal images. The vertical bars at the 2.0 and 3.0 sec mark indicated the duration of the image (1.0 sec) while the beach scene image was used to represent the general images and the dog image represented personal images. The red arcs of the brain map illustrated the topography of FC measures satisfying conditions p<0.01 (high FC values, red) and p>0.99 (low FC values, blue) at a point in time corresponding to peak values in NSFC plot. It should be noted that these FC changes occur in the person viewing the images (the Sender), as such these do not represent any evidence for anomalous interactions.

Significant FC increases and decreases in both cases were observed for participant Sender 1_1, although this effect appeared larger for the personal images when they appeared. Interestingly, prominent FC reductions prior to the appearance of the next image were observed for both the general and personal images. This may be suggestive of an anticipatory effect.

Each participant Receiver recording session yielded a result for the 50 general images and 50 personal images. Thus, in Session 1, the 10 participants yielded a set of 20 individual receiver findings (IRF). In Session 2, equipment malfunction resulted in 9 participants yielding a set of 18 IRFs. Thus, both sessions yielded a total of 38 individual receiver findings. Of the 38 individual receiver findings, 26 did not exhibit statistically significant FC changes according to the threshold criteria awhile the remaining 12 individual receiver findings exhibited at least one statistically significant NSFC peak.

Receiver 1_1 was one of three twins that exhibited significant NFSC values in both Sessions 1 and 2 (see Table 2 for summary). In Session 1, Receiver 1_1 exhibited a significant NSFC (24 significant FC measures, p=5X10^-3^) for the general images but not the personal images (not shown). In Session 2, Receiver R1_1 showed the largest NSFC (22 significant FC measures, p=4.3X10^-3^) for personal images and non-significant results for general images. In both cases, the effect was characterized by increases in Receiver Twin 1_1 occipito-parietal FC at the time that the images were being viewed by Sender Twin 1_2. The NSFC results for the control condition in Session 2, Fig 5B are shown as the dashed line which illustrates the positive or negative NSFC value, whichever is the larger number during the session 2 control condition when both twins are viewing a static image for 11 min. At no stage during of the control conditions did the number of significant FC measures (NSFC) reach statistical significance at the p≤0.01 level. None of the participants exhibited significant NSFC values during the Session 2 control condition.

**Fig 5A.**
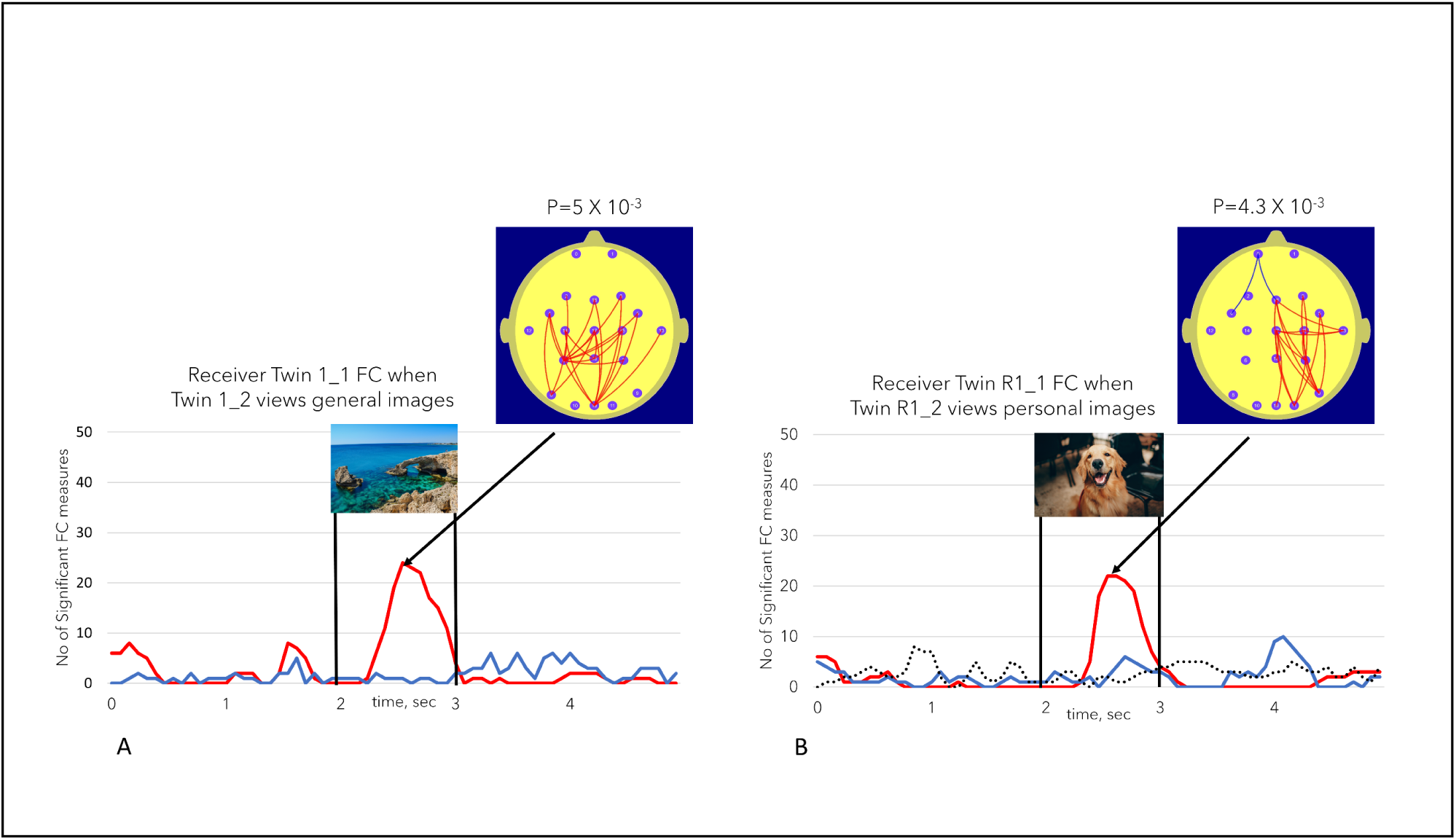
(left) illustrates the number of FC measures (NSFC) satisfying the conditions described in Fig 1 for <u>Receiver 1_1 FC</u> when sender 1_2 was viewing the general images. Fig 5B (right) illustrates the same <u>Receiver R1_1 FC</u> measures when sender R1_2 was viewing the personal images in session 2, or the repeat session. The dotted line in Fig 5B represents the NSFC measure during the session 2 control task when both twins were simultaneously viewing a static image for 11 min. The dotted line took the value of the maximum number of significant FC values (maxima or minima). In none of the session 2 control task sessions did the number of significant FC values reach the p<0.01 level.

**Table 1.**

|  | Pair 1 | Pair 2 | Pair 3 | Pair 4 | Pair 5 |
| --- | --- | --- | --- | --- | --- |
| Age | 21 years | 13 years | 27 years | 39 years | 17 years |
| MZ probability % | 99.9999998 | 99.99999999773 | 99.99999999766 | 99.99999999695 | 99.99999999339 |

**Table 2.**
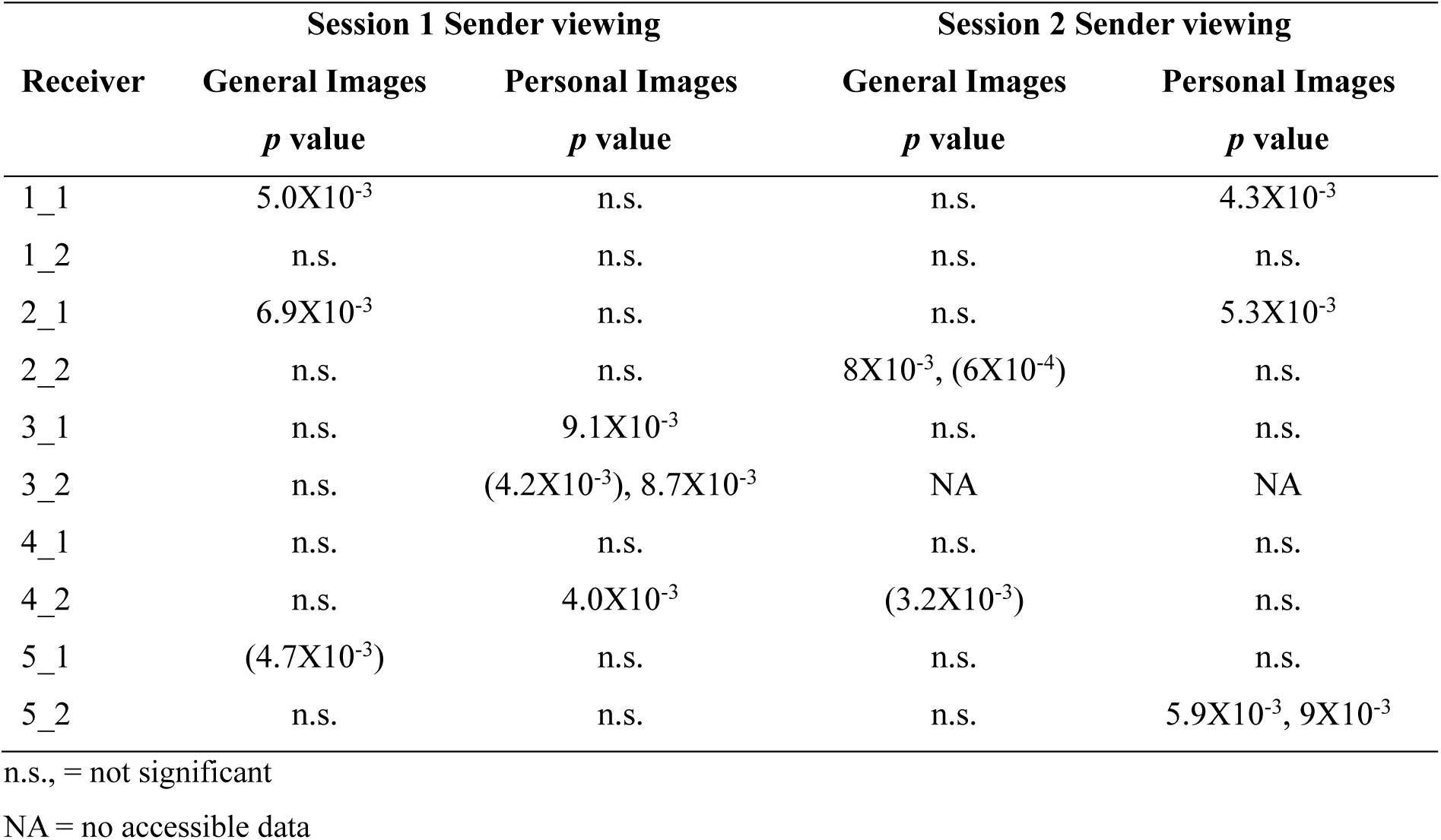

The patterns of FC changes demonstrated large inter-individual variation. Fig 6 illustrates another of the statistically significant individual Receiver findings. Fig 6A illustrates FC changes in Receiver T3_2 during session 1 when sender T3_1 was viewing personal images in this session. In this case, the early FC decreases were primarily located at left temporal to prefrontal sites while the later FC increases exhibited bilateral temporal to prefrontal increases. By contrast, Receiver T2_2 in session 2 exhibited an early FC increase with a left parieto-temporal to frontal and central sites with a later more extensive bilateral FC decrease. The dotted line in Fig 5B once again illustrates the number of significant FC changes (maximum of either positive or negative) during the session 2 control condition when both twins are viewing a static image for 11 min.

**Figs 6A and 6B.**
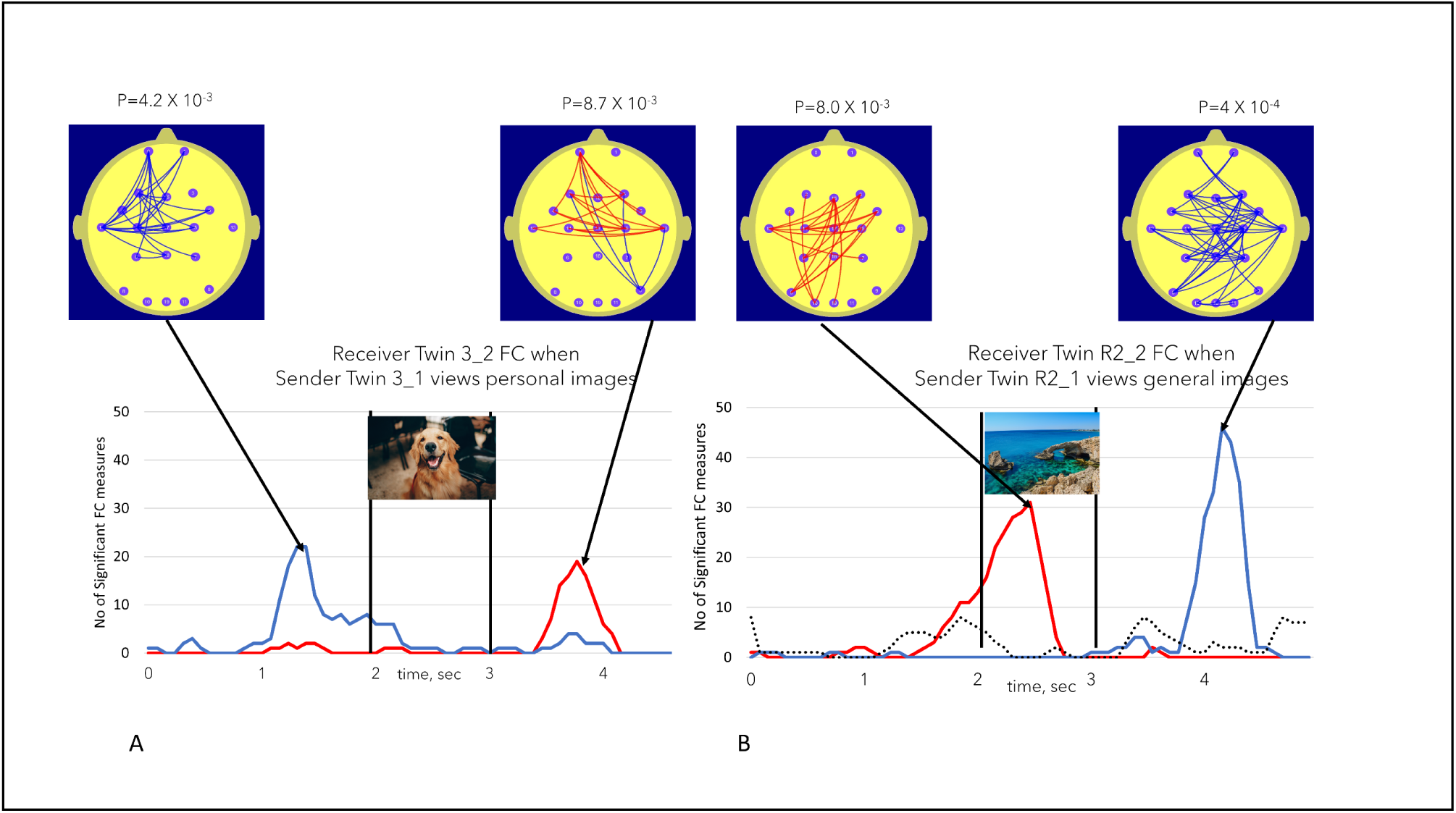
illustrate the variations in FC increase or decrease, FC topography and the timing of these FC changes in two participants. Fig 6A illustrates significant findings in Receiver 3_1 when Sender 3_2 was viewing personal images. Fig 6B illustrates the equivalent finding for Receiver 2_2 during session 2 or the repeat session ie R2_2 when Sender 2_1 is viewing general images. The dotted line in Fig 6B represents the NSFC measure during the session 2 control task when both twins were simultaneously viewing a static image for 11 min. The dotted line took the value of the maximum number of significant FC values (maxima or minima). In none of the session 2 control task sessions did the number of significant FC values reach the p<0.01 level.

Table 2. summarizes the uncorrected probability of observing the statistically significant NSFC findings for sessions 1 and 2.

Findings listed in brackets refer to significant effects due to FC decreases while the others refer to corresponding FC increases.

To summarize, in the 38 IRFs listed in Table 2, we observed 7 cases where NSFC values were associated a probability in the range of 0.005≤p<0.01 and 7 cases where NSFC values were associated with a probability in the range p<u><0</u>.005. To determine the experiment wide statistical significance of these findings, the trinomial distribution was used. To determine the overall number of degrees of freedom we note that each individual 5.0 sec epoch is associated with 4.6 degrees of freedom. With 38 individual receiver findings and 4.6 degrees of freedom per individual receiver finding, the experiment wide number of degrees of freedom was estimated as 38 X 4.6 or 174.8 rounded up to 175.

The trinomial distribution yielded an overall p=4 X 10^-8^ with Df=175. Calculating the experimental effect size, Cohen’s d equivalent (d_equ) using the approach described by Rosenthal & Rubin (2003) yielded the following:

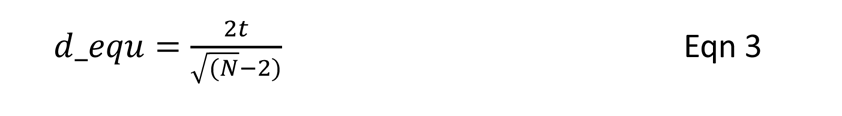

In equation 3, t is an equivalent one tailed student’s t value based on the calculate p value (p=4X10^-8^) and N=175 degrees of freedom. We used the Excel T.INV function to determine the t-value for the nominated p and N values. This yielded a value of t=5.607 and a resulting Cohen’s d equivalent d_equ=0.85 which is considered a large experimental effect size (Cohen 2013).

We observed 8 statistically significant NSFC findings when the Sender was viewing personal images and 6 when the Sender was viewing general images and 6 equivalent cases where the Sender was viewing general images. A binomial distribution indicated this difference was not statistically significant p=0.2 Df=14.

While there were large inter-individual differences in the Receiver FC topography in the statistically significant individual findings, the following briefly summarizes the main topographic FC features we observed.

Of the 14 significant Receiver findings, 10 were associated with FC increases and 4 FC decreases. A binomial distribution indicated that this difference approached but did not reach statistical significance, p=0.06, Df=14. In terms of hemispheric differences, 6 statistically significant individual FC findings were located predominantly in the left hemisphere, 5 extended bilaterally, while 3 were located in the right hemisphere. When considering the topography of the individual FC findings, the following regions most frequently involved (and listed in descending order of occurrence) were parieto-temporal, frontal-prefrontal and occipito-parietal.

## Discussion

To the best of the author’s knowledge, this is the first study reporting dynamic changes in brain FC associated with anomalous interactions between monozygotic twins. Specifically, we observed statistically significant changes in receiver FC when the event related FC measure was determined using the sender viewing times. As such Hypothesis 1 was confirmed.

The findings are thus consistent with the studies reviewed in the introduction that report correlated biometric or brain activity between pairs of individuals that share a close emotional relationship. Of particular note is the fMRI study reporting anomalous interactions in a set of monozygotic twins (Karavasilis et al 2018). Apart from the Duane & Behrendt (1965) paper, these were the only studies we could identify that reported biometric or brain activity changes suggesting anomalous interactions between identical twins.

While we found that there were more statistically significant cases when Senders were viewing personal images (8) compared to general images (6), the small numbers made this difference statistically insignificant. Thus, hypothesis 2 was not confirmed.

A prominent feature of the results was the large intersubject variability. While only approximately 1/3 of the individual findings were associated with statistically significant FC increases or decreases, these FC features exhibited significant inter-individual variability in terms of FC topography or regions involved in the FC changes and the timing of the receiver FC changes. It should be noted that while FC increases and decreases were observed in the Receiver data, such FC increases and decreases were also observed in the sender data as illustrated in Fig 3. Furthermore, such FC increases and decreases and have also been observed during cognitive tasks reported in other studies (Bressler & Kelso 2016, Silberstein & Camfield 2021). We have generally interpreted such FC increases and decreases as an indication of cortical information processing where increased FC represents stronger information transfer between networks and reduced FC as an inhibition of network communication which may interfere with the efficiency of cortical processing (Silberstein et al 2017).

While relatively large individual differences in regional brain activity and functional connectivity are commonly observed in cognitive neuroscience brain imaging studies (Kania & Rees 2011), these findings suggest an additional layer of complexity in interpreting individual receiver data based on what the sender is viewing. Specifically, the receiver FC changes are presumed to be associated with the receiver’s unconscious subjective responses which are, in turn, presumed to be influenced by the sender’s subjective response to the images being presented. In other words, what we observe as Receiver FC changes may be influenced by two successive layers of subjective response. Each of these two layers of subjective responses likely add a significant degree of inter-individual variability. Thus, expecting a consistent relationship between the content of the images as viewed by the Sender and the Receiver FC changes is unrealistic. More generally, if such large inter-individual variability is a common feature of FC changes associated with anomalous interactions, then this suggests that simple pooling or cross-subject averaging is likely to dilute individual effects and yield non-significant findings.

Another factor warranting comment is the relatively strong experimental effect size observed (Cohen’s d=0.85). This is a relatively high effect size, especially for studies examining anomalous cognition effects (Cardena 2018). It is possible that several experimental factors contributed to this high effect size. One factor may be the recruitment method. By limiting recruitment to monozygotic twins, the chance of observing experimental statistically significant effects may have increased. Past research has shown that monozygotic twins tend to report stronger emotional attachment with their twin (Fortuna et al 2010, Landenberger et al 2021) and a higher prevalence of anomalous shared experiences (Brusewitz et al 2013), when compared to fraternal (dizygotic; DZ) twins. Additionally, participants were required to have previously shared an anomalous experience. This may have also contributed as past studies have shown that a belief in the reality of anomalous cognition effects is associated with a greater likelihood of exhibiting positive findings (Lawrence, 1993).

Another factor is the choice of Steady State Visually Evoked Potential Event Related Partial Coherence (SSVEP-ERPC) as the methodology used to measure FC changes. This methodology has been used to examine FC and FC changes in areas as diverse as the neural correlates of mental rotation (Silberstein et al 2003), creativity (Silberstein et al 2019, 2021) and attention deficit hyperactivity disorder (Silberstein et al 2016A, 2016B, 2017). Previous research has found this methodology to be especially sensitive to the FC correlates of cognitive processes. These processes are now thought to involve top-down or feedback communication between cortical neural networks and are understood to be mediated by synchronous oscillations in the 10Hz – 20Hz range (Bressler & Richter 2015, Fries 2015). As this methodology is only sensitive to the synchronous oscillations or FC occurring at the stimulus frequency of 13Hz, this means that top-down processes are primarily registered, and these may be the ones that play a significant role in processing the anomalous interactions observed.

Finally, the relatively high effect size may, in part, be due to the visual flicker stimulus frequency. This is the fact that both twins were exposed to the same 13Hz stimulus used to elicit the SSVEP. In other words, both stimuli were precisely synchronized with a zero-phase difference. Whilst highly speculative, this factor is based on findings emerging from EEG hyper-scanning studies where positive social interactions between individuals was associated with increased intersubject synchronization of EEG power in various frequency ranges (Muller et al 2022). An intriguing finding suggesting that eliciting interbrain synchronization may enhance positive social interactions was reported by Yang et al (2021). In this study, rat frontal cortex was activated opto-genetically using an implanted light source. The authors report that the effect of opto-genetically stimulating two rats at the same frequency is to increase positive social interactions between the rats. This occurs even if the rats exhibit mutual antagonistic behaviour before the optogenetic stimulation. Taking account of the human hyper scanning studies pointing to synchronous brain oscillations increased positive social interactions together with the intriguing findings reported by Yang et al (2021), it may be that exposing both the sender and receiver to the same visual flicker stimulus frequency may enhance the strength of the anomalous interaction. The current experimental design cannot determine whether this hypothetical ‘interbrain synchrony’ effect was present or not. However, future studies will explore the sensitivity of the anomalous interaction FC effects to differences in sender-receiver flicker frequency.

Given the nature of the findings, it is important that this study be independently replicated. Furthermore, any such a replication should be consistent with certain key features of the present study. Firstly, it is important to recruit monozygotic twins who report having experienced anomalous interaction and are therefore positively inclined to believe these effects are real. In addition, it is important to use the SSVEP-ERPC methodology with a 13Hz flicker frequency. Given that the FC components observed using the SSVEP-ERPC were primarily a reflection of top-down effects, the specific features of the methodology are considered important factors contributing to the experimental size effect.

Typically, the discussion section of a paper describing an experimental finding will include a consideration of the reported findings in the context of previous research and the extent to which the new findings confirm or challenge established models and understandings. However, the subject matter of the current findings makes this difficult as there is as yet no widely accepted model that can be used to interpret our findings in terms of currently understood physical or biological processes. Some researchers have pointed to quantum mechanics as an approach that may provide a theoretical model for the type of findings we have reported. For example, the phenomenon of quantum entanglement has been suggested variously as either a metaphor or as the basis of a physical model for the type of findings we and others have reported (Cardena 2018, Kauffman & Radin 2023, Radin 2009, Tressoldi et al 2010). Furthermore, a series of replicated findings indicating a link between conscious intent and changes in the double slit photon interference pattern reinforces the notion of a possible link between quantum processes and the type of anomalous phenomena we are reporting (Radin et al 2012).

While quantum entanglement may indeed provide a useful metaphor and perhaps even a model for the non-local nature of our observations, we take an agnostic position in this regard. While we cannot point to a widely accepted physical model to interpret our findings, we consider the report of such experimental findings a necessary step in the eventual development of such a model. The scientific method is based on a foundation of empiricism where novel veridical observations come first and a theory or model to account for the observations is then developed. If our observations are correct and independently replicated, then they, in conjunction with the large body of comparable replicated evidence (Cardena 2018) may contribute to the development of a deeper and more comprehensive model capable of accounting for such phenomena.

### Limitations and Future Directions

Finally, this study suffers from several limitations. Although 10 participants make this one of the larger studies its kind, the number of participants is too small to reliably identify any consistent features in the topography or timing of the FC components. It is therefore recommended that future studies recruit a larger number of participants. Another important limitation is the relatively small number of scalp recording sites. The small number of electrodes severely reduces the spatial resolution of the findings and makes it impossible to use inverse mapping techniques that would enhance the ability to accurately identity which cortical areas are contributing to the FC changes. Future studies could employ 64 electrode headsets, thus greatly improving spatial resolution. Another limitation stemmed from the use of a 20m cable to ensure that both twins were exposed to the same 13Hz stimulus. This meant that twins could not be spatially separated by more than 20m. While this is not a serious limitation, it hinders the ability to explore the effect of spatial separation on the anomalous interactions. Subsequently, recording systems capable of precisely synchronizing the flicker stimulus between twins without the necessity of using a cable would allow spatial separation to be explored.

## Conclusion

Results from this study support the presence of anomalous interactions between sensorily isolated monozygotic twins. Specifically, statistically significant changes were observed in Receiver FC when trials were synchronized to the Sender image. The novel findings of this study, in conjunction with the unique application of methodology, necessitates future replication and further research.

## References

Acharya, J. N., Hani, A. J., Cheek, J., Thirumala, P., and Tsuchida, T. N. (2016). American Clinical Neurophysiology Society guideline 2: guidelines for standard electrode position nomenclature. Neurodiagn. J. 56, 245–252. doi: 10.1080/21646821.2016.1245558

Achterberg, J., Cooke, K., Richards, T., Standish, L. J., Kozak, L., & Lake, J. (2005). Evidence for correlations between distant intentionality and brain function in recipients: A functional magnetic resonance imaging analysis. *Journal of Alternative & Complementary Medicine: Research on Paradigm*, Practice, and Policy, 11(6), 965–971

Ambach, W. (2009). Correlations between the EEGs of two spatially separated subjects—a replication study. European Journal of Parapsychology, in print.

Andrew, C., & Pfurtscheller, G. (1996). Event-related coherence as a tool for studying dynamic interaction of brain regions. Electroencephalography and clinical neurophysiology, 98(2), 144–148.

Billows, H., & Storm, L. (2016). Believe it or not: A confirmatory study on predictors of paranormal belief, and a psi test. Australian Journal of Parapsychology, 15(1), 7–35.

Bouvet, R., & Bonnefon, J. F. (2015). Non-reflective thinkers are predisposed to attribute supernatural causation to uncanny experiences. Personality and Social Psychology Bulletin, 41(7), 955–961.

Bressler, S. L., & Kelso, J. S. (2016). Coordination dynamics in cognitive neuroscience. Frontiers in Neuroscience, 10, 397.

Bressler, S. L., & Richter, C. G. (2015). Interareal oscillatory synchronization in top-down neocortical processing. Current opinion in neurobiology, 31, 62–66.

Brusewitz, G., Cherkas, L., Harris, J., & Parker, A. (2013). Exceptional experiences amongst twins. J. Soc. Psychical Res., 1–14.

Cardeña, E. (2018). The experimental evidence for parapsychological phenomena: A review. Am. Psychol., 73(5), 663–677. 10.1037/amp0000236

Cohen, J. (2013). Statistical power analysis for the behavioral sciences. Academic press.

Duane, T., & Behrendt, T. (1965). Extrasensory electroencephalographic induction between identical twins. Science, New Series, 150(3694).

Fortuna, K., Goldner, I., & Knafo, A. (2010) Twin relationships: A comparison across monozygotic twins, dizygotic twins, and nontwin siblings in early childhood. Family Sci., 1, 205–211. 10.1080/19424620.2010.569367

Fries, P. (2015). Rhythms for Cognition: Communication through Coherence. Neuron, 88, 220–235.

Gray M, Kemp AH, Silberstein RB, Nathan PJ (2003) Cortical neurophysiology of anticipatory anxiety: an investigation utilizing steady state probe topography (SSPT). Neuroimage. 20:975–986.

Kanai, R., & Rees, G. (2011). The structural basis of inter-individual differences in human behaviour and cognition. Nature Reviews Neuroscience, 12(4), 231–242.

Karavasilis, E., Christidi, F., Platoni, K., Ferentinos, P., Kelekis, N. L., & Efstathopoulos, E. P. (2018). Functional MRI study to examine possible emotional connectedness in identical twins: A case study. Explore, 14(1), 86–91.

Kauffman, S. A., & Radin, D. (2023). Quantum aspects of the brain-mind relationship: A hypothesis with supporting evidence. Biosystems, 223, 104820.

Landenberger, R. D. O., Lucci, T. K., David, V. F., França, I., Souza, E. De, Segal, N., Otta, E. (2021). Hierarchy of attachment figures among adult twins and non-twins. Pers. Individ. Differ., 170, 110404. 10.1016/j.paid.2020.110404

Lawrence, T. (1993). Gathering in the sheep and goats: A meta-analysis of forced-choice sheep/goat ESP studies, 1947-1993. Proceedings of the Parapsychological Association 36th Annual Convention, Toronto, Canada, 75-86.

Moulton, S. T., & Kosslyn, S. M. (2008). Using neuroimaging to resolve the psi debate. Journal of Cognitive Neuroscience, 20(1), 182–192.

Müller, V., Fairhurst, M. T., van Vugt, F. T., Keller, P. E., & Müller, M. F. (2022). Interpersonal synchrony and network dynamics in social interaction. Frontiers in Human Neuroscience, 16, 1095735.

Panofsky, H.A. and Brier, G.W., 1958. Some applications of statistics to meteorology. Mineral Industries Extension Services, College of Mineral Industries, Pennsylvania State University.

Parker, A., & Jensen, C. (2013). Further possible physiological connectedness between identical twins: The London study. Explore, 9(1), 26–31.

Radin, D. I. (2004). Event-related electroencephalographic correlations between isolated human subjects. The Journal of Alternative & Complementary Medicine, 10(2), 315–323.

Radin, D. (2009). Entangled minds: Extrasensory experiences in a quantum reality. Simon and Schuster.

Radin, D. (2017). Electrocortical correlations between pairs of isolated people: A reanalysis. F1000Research, 6(676). 10.12688/f1000research.11537.1

Radin, D., Michel, L., Galdamez, K., Wendland, P., Rickenbach, R., & Delorme, A. (2012). Consciousness and the double-slit interference pattern: Six experiments. Physics Essays, 25(2).

Radin, D. I., & Schlitz, M. J. (2005). Gut feelings, intuition, and emotions: An exploratory study. Journal of Alternative & Complementary Medicine, 11(1), 85–91.

Rosenthal, R., & Rubin, D. B. (2003). R equivalent: A simple effect size indicator. Psychological methods, 8(4), 492.

Maris, E., Schoffelen, J. M., & Fries, P. (2007). Nonparametric statistical testing of coherence differences. Journal of neuroscience methods, 163(1), 161–175.

Silberstein R B, Danieli F, Nunez PL. (2003) Fronto-parietal evoked potential synchronization is increased during mental rotation. Neuroreport. 14:67–71.

Silberstein R.B, Camfield D.A, Nield G, Stough C. (2019) Gender differences in parieto-frontal brain functional connectivity correlates of creativity. Brain Behav. 2019;9:e01196. 10.1002/brb3.1196

Silberstein, R. B., & Camfield, D. A. (2021). Sex influences the brain functional connectivity correlates of originality. Scientific reports, 11(1), 1–10.

Silberstein, R. B., Pipingas, A., Farrow, M., Levy, F., Stough, C. K., & Camfield, D. A. (2016A). Brain functional connectivity abnormalities in attention-deficit hyperactivity disorder. Brain and behavior, 6(12), e00583.

Silberstein, R. B., Pipingas, A., Farrow, M., Levy, F., & Stough, C. K. (2016B). Dopaminergic modulation of default mode network brain functional connectivity in attention deficit hyperactivity disorder. Brain and behavior, 6(12), e00582.

Silberstein, R. B., Levy, F., Pipingas, A., & Farrow, M. (2017). First-dose methylphenidate– induced changes in brain functional connectivity are correlated with 3-month attention-deficit/hyperactivity disorder symptom response. Biological psychiatry, 82(9), 679–686.

Standish, L. J., Johnson, L. C., Kozak, L., & Richards, T. (2003). Evidence of correlated functional magnetic resonance imaging signals between distant human brains. Alternative Therapies in Health and Medicine, 9(1), 128–128.

Standish, L. J., Kozak, L., Johnson, L. C., & Richards, T. (2004). Electroencephalographic evidence of correlated event-related signals between the brains of spatially and sensory isolated human subjects. The Journal of Alternative & Complementary Medicine, 10(2), 307–314

Storm, L., & Tressoldi, P. E. (2017). Gathering in more sheep and goats: A meta-analysis of forced-choice sheep-goat studies, 1994-2015. Journal of the Society for Psychical Research, 81(2), 79-107.

Tressoldi, P. E., Storm, L. & Radin, D. (2010). Extrasensory perception and quantum models of cognition. NeuroQuantology, 8(4), 581–587.

Wackermann, J. (2004). Dyadic correlations between brain functional states: present facts and future perspectives. Mind and Matter, 2(1), 105–122.

Wackermann, J., Seiter, C., Keibel, H., & Walach, H. (2003). Correlations between brain electrical activities of two spatially separated human subjects. NeuroScience letters, 336(1), 60–64.

Yang, Y., Wu, M., Vázquez-Guardado, A., Wegener, A. J., Grajales-Reyes, J. G., Deng, Y., … & Rogers, J. A. (2021). Wireless multilateral devices for optogenetic studies of individual and social behaviors (pp. 1-11). Nature Publishing Group.

Yu, J.-Y. (2005) ESS210B Lectures, Time Series. https://www.ess.uci.edu/~yu/class/ess210b/lecture.4.time.series.all.pdf

